# GJB3, a gap junction gene, supports cell growth by mediating cystine uptake and regulating cellular stress pathways in SLC7A11 low adenocarcinomas

**DOI:** 10.1101/2024.12.03.626548

**Authors:** Disha Acharya, Annesha Chatterjee, Nikita Bhandari, Payel Roy, Monishka Agrawal, Bal Krishna Chaube, Sudhanshu Shukla

**Author notes:** Contributed equally (Bal Krishna Chaube, Sudhanshu Shukla).

## Abstract

Gap junctions are specialized intercellular connections that directly connect the cytoplasm of two cells via protein structure called connexins. Despite extensive research on many cell surface proteins, Gap junction proteins are understudied in cancer. In this study, we used TCGA data to identify genetic and epigenetic changes associated with Gap junction proteins. The analysis identified GJB3 as a key gene with differential methylation and expression patterns, with notable overexpression in COAD and LUAD, correlating significantly with patient survival outcomes. GJB3 knockdown studies revealed reduced cell proliferation and migration. Transcriptomic analysis revealed that GJB3 knockdown induced a cellular stress response, characterized by activation of starvation and autophagy pathways. Western blot analysis confirmed these findings, showing increased phosphorylation of eIF2α and activation of the GCN2-eIF2α-ATF4 signaling axis subsequent autophagy induction. We also found that sustained autophagy induced apoptosis mediated cell death. Metabolic profiling revealed a significant decrease in cystine levels in GJB3-deficient cells. We demonstrated that GJB3 plays a crucial role in cystine uptake, especially in cells with low SLC7A11 expression. Furthermore, we showed that GJB3 can be targeted using specific antibodies, establishing it as a potential therapeutic strategy for GJB3-dependent cancers. These findings highlight the significance of GJB3 in cancer progression and its potential as a therapeutic target, offering new insights into its epigenetic regulation and functional role in cellular stress and survival mechanisms.

## 2. Introduction

Cell surface proteins (CSPs) play a crucial role in cancer biology, acting as both biomarkers for disease progression and targets for therapeutic interventions (1). These proteins are essential for various cellular processes, including cell signalling, adhesion, and modulation of the immune response (2). The cancer cell surfaceome has emerged as a promising target for developing targeted therapies, particularly with proteins such as EGFR, which is frequently overexpressed in many solid tumors and is the focus of monoclonal antibody therapies and tyrosine kinase inhibitors (3). While CSPs like EGFR have been extensively studied, gap junctions remain relatively underexplored in the context of cancer (4).

Gap junctions are specialized intercellular channels formed by connexin proteins that enable direct communication between adjacent cells (5). They facilitate the exchange of ions, small molecules, and signalling molecules, playing a vital role in physiological processes such as cardiac electrical activity, tissue synchronization, and embryonic development (6,7). Historically, connexins were thought to function primarily as tumor suppressors by enhancing cell-to-cell communication (4). Cx43 (coded by GJA1) is the most studied Gap junction protein in cancer (8). Many studies have correlated the expression of Cx43 with improved patient prognosis in prostate cancer, breast cancer, non-small cell lung cancer and colorectal cancer (9–12). However, contradictory evidence exists; for instance, Cx43 expression has been associated with poor survival outcomes in oral, esophageal, and bladder cancers (13–15). Similar conflicting associations have been observed for other gap junction proteins such as Cx26 and Cx30, depending on the specific cancer type (4).

Functionally, gap junction proteins exhibit variable phenotypes across different cancers. For example, Cx43 has been shown to suppress the EMT and reduce the cisplatin chemoresistance of A549 cells. However, Cx43 helps in promoting brain metastasis by cGAMP transfer between carcinoma and astrocytes (16). Similarly, Cx32 has been linked to increase in cancer stem cells self-renewal in human liver cancer cells (17). Furthermore, studies indicate that the expression of gap junction proteins correlates with various hallmarks of cancer, including epithelial-mesenchymal transition (EMT), differentiation, and angiogenesis (16,18,19). These observations underscore the complex roles of gap junction proteins in cancer biology.

Recent findings highlight that GJC1 is hypermethylated and silenced in colorectal cancer and related cell lines (20). Additionally, Cx43 regulation by microRNAs such as miR-221/222 in glioma and miR-20A in prostate cancer has been documented (21,22). While some gap junction proteins have been implicated in tumor suppression, emerging evidence suggests a more nuanced role. For instance, GJB3 has been downregulated in bladder cancer, contributing to aneuploidy (23); conversely, Cx26 promotes tumor progression by regulating metastatic potential through pathways like PI3K-AKT and conferring resistance to EGFR inhibitors (24). Despite numerous studies detailing the functions of GJB3, many other gap junction proteins remain underexplored within cancer research. Our research identifies a novel role for GJB3 in lung and colorectal cancers, presenting it as a hypomethylated and overexpressed gene associated with poor patient prognosis. We demonstrate that GJB3 knockdown inhibits cell growth, induces a cellular starvation response, and is crucial for cystine transport in cells with low SLC7A11 expression. Additionally, we show that GJB3 can be targeted therapeutically using a specific antibody. This underscores the potential of gap junction proteins not only as biomarkers but also as therapeutic targets in oncology.

## 3. Results

### 3.1 GJB3 is regulated by methylation in four common adenocarcinomas

To understand the genetic aberration associated with Gap junction genes, we used mutation and copy number aberration data from cBioPortal. The analysis showed that none of the gap junction proteins showed high mutation rate which are oncogenic in nature (**Figure 1A**). We noticed GJA3 (Amplified in COAD), GJA5 (Amplified in LUAD and BRCA), GJA8 (Amplified in LUAD and BRCA), GJA10 (Deleted in PRAD), GJB7 (deleted in PRAD), and GJC2 (Deleted in BRCA) show copy number variation in more than 5% of samples of individual cancers (**Supplementary figure 1A**). Next, we checked if CNV is regulating the gene expression of these amplified genes. Interestingly, GJA3, GJA8, GJA10 and GJB7 are not expressed in any of the four cancers or corresponding normal and other two genes did not show any correlation with expression and CNV (**Figure 1B and Supplementary figure 1B and C**), suggesting these genes are not regulated by CNV in these cancers. We also checked the expression of these genes in four cancer types and corresponding normal. Only nine Gap junction proteins (shown in figure 1B with arrow) are expressed in these four cancers and corresponding normal. Given the limited understanding of GJB3’s role in cancer, we decided to study this gene in further detail to explore its potential implications in cancer progression and its significance as a therapeutic target.

**Figure 1:**
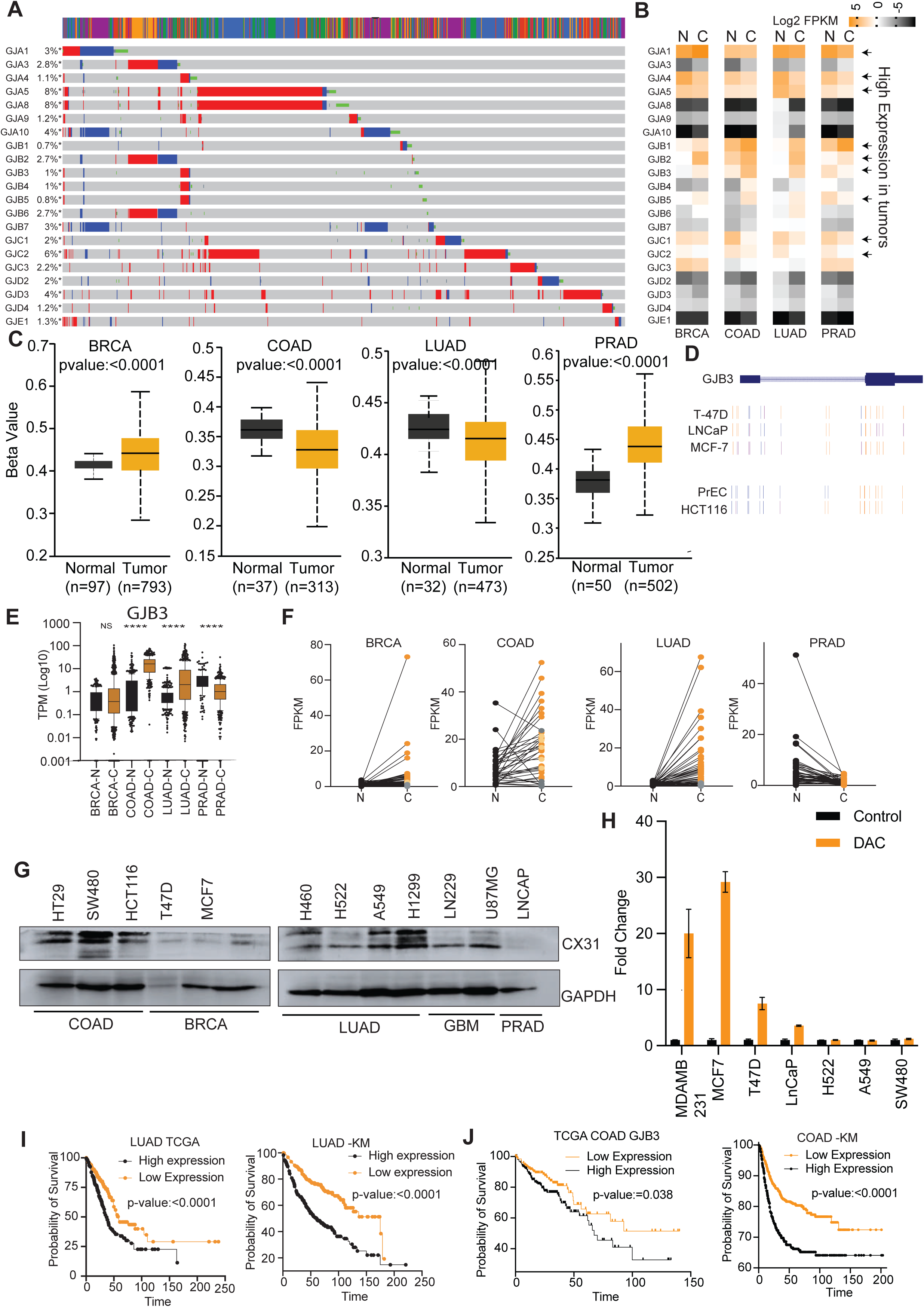
GJB3 regulated by DNA hypomethylation: **A)** Genetics data for Gap junction genes was analyzed using cBioPortal. The Oncoprint plot for four cancer types are shown. **B)** The expression of all the 21 Gap junction genes was analyzed in cancer and normal samples and. The genes with >1 FPKM were considered expressed genes. The arrows indicate the genes with good expression in cancer. **C)** The beta values of CpGs associated with GJB3 in normal and cancer tissues of BRCA, COAD, LUAD and PRAD was plotted in UALCAN data base and shown here. **D)** The methylation values of CpGs associated with the GJB3 gene in different cell lines from breast, prostate, and colon cancers were obtained from the UCSC Genome Browser and displayed. **E)** GJB3 expression levels were compared in four adenocarcinoma samples and their corresponding normal tissues. The results are plotted with p-values indicated as **** <0.0001. **F)** Comparison of GJB3 expression between four adenocarcinoma samples and their matched normal tissues. **G)** The expression of the GJB3 gene in various cell lines was measured using western blot. GAPDH was used as internal control. **H)** Selected cells were treated with a given amount of deoxy-azacytidine (DAC), and expression was measured using q-RT PCR. The fold changes compared to control were measured and plotted. **I)** GJB3 expression was correlated with survival rates in lung adenocarcinoma (using TCGA and KM plotter data) and KM Plots were developed and shown. **J)** GJB3 expression was correlated with survival rates in colorectal cancer (using TCGA and KM plotter data). and KM Plots were developed and shown.

To understand the mechanism of GJB3 deregulation in four adenocarcinomas, we checked the DNA methylation of the CpGs of the GJB3 promoter using TCGA data from the UALCAN database. We found that GJB3 promoter was significantly hypermethylated in BRCA and PRAD and significantly hypomethylated in COAD and LUAD compared to corresponding normal samples (**Figure 1C**). To confirm the methylation-mediated regulation of GJB3, we utilized the UCSC genome browser and methylation data associated with various cell lines. This approach allowed us to validate the observed patterns of GJB3 regulation by DNA methylation in cell line models (**Figure 1D**).

We investigated GJB3 expression levels in four adenocarcinomas: COAD, LUAD, PRAD, and BRCA. Analysis revealed increased GJB3 expression in colon and lung cancers, while breast cancer showed no significant difference compared to healthy tissue. Prostate cancer displayed decreased GJB3 expression (**Figure 1E**). This pattern held true when analyzing matched tumor and normal tissue samples (**Figure 1F**).

Next, we assessed the expression of Cx31 (protein coded by GJB3 gene) in various cancer cell lines and found that it was expressed in all colorectal cancer cell lines and some lung cancer cell lines (**Figure 1G**). This observation aligned with the decreased methylation and increased expression of GJB3 in COAD and LUAD tumor samples. To further confirm the methylation-mediated regulation of GJB3we treated cells with 5-Deoxy-Azacytidine (DAC), a DNA methyltransferase inhibitor and check the expression of GJB3 RNA. Our results showed that in cells where GJB3 was not expressed, DAC treatment led to an increase in its expression (**Figure 1H**). Conversely, cells with high GJB3 expression, such as SW480, did not exhibit a significant effect on GJB3 levels upon DAC treatment (**Figure 1H**).

Next, we explored the potential role of GJB3 in patient survival for lung and colon adenocarcinomas (**Figure 1I and J**). GJB3 expression emerged as a significant predictor of survival in both cancer types, based on data from two independent patient cohorts (TCGA and KM plotter) (**Figure 1I and J**).

These findings collectively support the notion that DNA methylation plays a crucial role in regulating GJB3 expression in colorectal and lung adenocarcinomas.

### 3.2 GJB3 is overexpressed in colorectal and lung cancer and predicts patients’ outcome

To understand GJB3’s cellular function in cancer, we employed knockdown studies. Using two shRNAs, we successfully reduced GJB3 levels in cancer cells, which was confirmed through PCR (**Figure 2A**). Interestingly, we found that GJB3 knockdown cells showed decreased cell proliferation and decreased colony formation activity (**Figure 2B and C**). We also measured the effect of GJB3 knockdown on cell migration and showed that GJB3 knockdown cells migrate significantly slower than control cells (**Figure 2D)**. Additionally, imaging analysis revealed increased cell death in GJB3 knockdown cells (**Figure 2E**).

**Figure 2:**
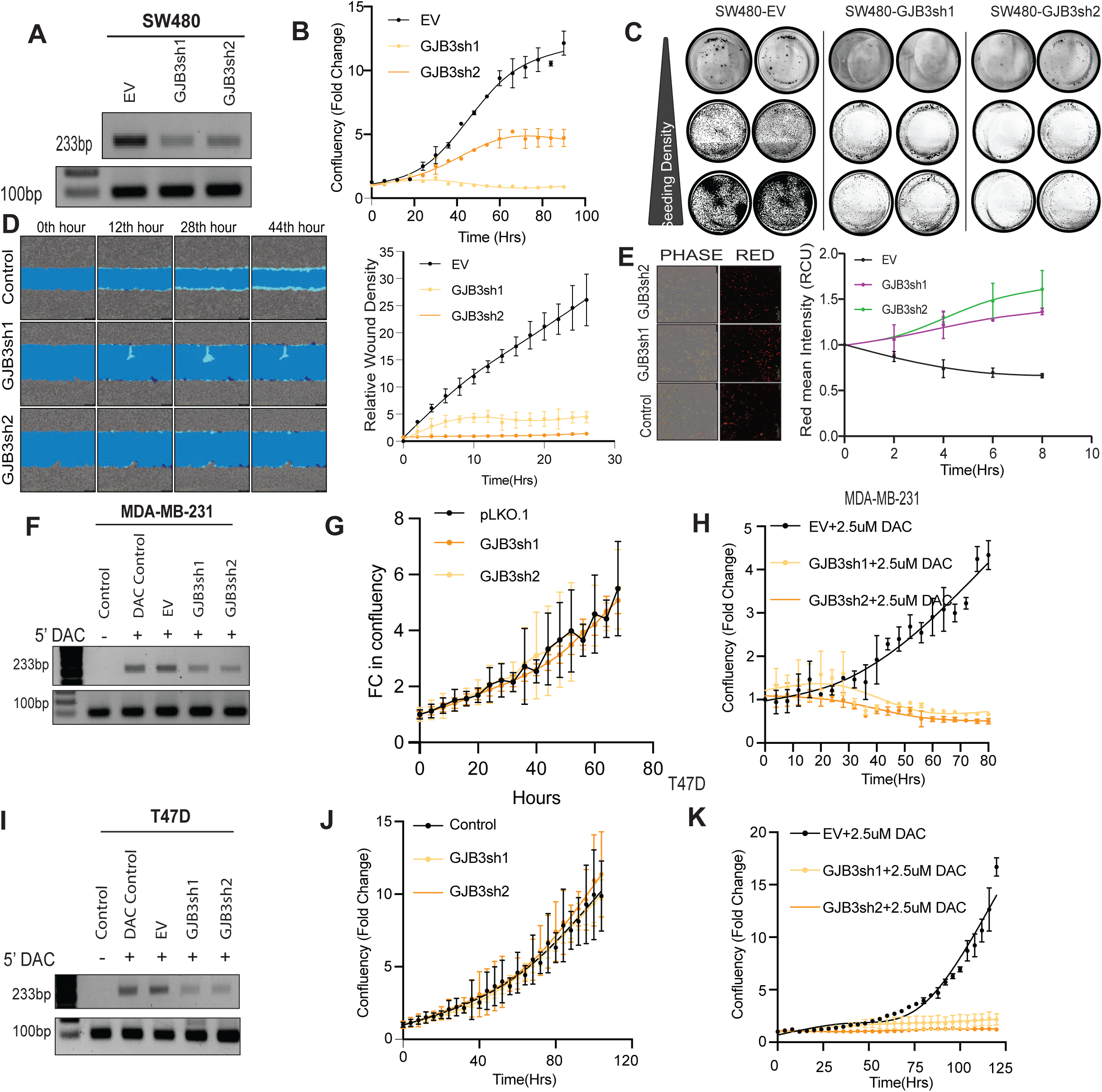
GJB3 is required for the growth in some adenocarcinoma cells. **A)** SW480 cells were subjected to GJB3 knockdown using two shRNAs, and the expression was measured by semi-quantitative PCR, with TPT1 used as a control. **B)** Proliferation assays were conducted on SW480 GJB3 knockdown and control cells using the Incucyte live cell imaging system. Fold change in confluency was plotted over 0-96 hours. **C)** Colony suppression assays were performed on SW480 GJB3 knockdown and control cells. Cells were seeded at three different concentrations and incubated for 15 days, followed by washing and staining with Coomassie blue. **D)** Wound healing assays were conducted on SW480 GJB3 knockdown and control cells. After a 12-hour incubation period, a scratch was made, and cell migration was measured using the Incucyte wound healing module. **E)** SW480 GJB3 knockdown and control cells were stained with propidium iodide, and images were captured at various time points using the Incucyte live cell imaging system. **F)** MDA-MB231 cells were infected with control or sh1 and sh2 GJB3 shRNA, treated with 10 μM DAC, and RNA was isolated. GJB3 expression was analyzed using semi-quantitative PCR and agarose gel electrophoresis, with TPT1 as a control. **G)** Proliferation of MDA-MB231 cells infected with control or sh1 and sh2 GJB3 shRNA was measured using the Incucyte live imaging system. **H)** MDA-MB231 cells infected with control or sh1 and sh2 GJB3 shRNA were treated with 10 μM DAC, and proliferation was measured using the Incucyte live imaging system. **I)** T47D cells were infected with control or sh1 and sh2 GJB3 shRNA, treated with 10 μM DAC, and RNA was isolated. GJB3 expression was analyzed using semi-quantitative PCR and agarose gel electrophoresis, with TPT1 as a control. **J)** Proliferation of T47D cells infected with control or sh1 and sh2 GJB3 shRNA was measured using the Incucyte live imaging system. **K)** T47D cells infected with control or sh1 and sh2 GJB3 shRNA were treated with 10 μM DAC, and proliferation was measured using the Incucyte live imaging system

To understand how GJB3 expression affects response to demethylating agents, we focused on breast cancer cells (MDA-MB231 and T47D) where GJB3 expression might be suppressed by hypermethylation. We treated MDAMB231 cells with a demethylating agent (DAC) at a concentration of 2.5μM, which led to increased GJB3 RNA levels (**Figure 2F**). Next, we infected DAC treated MDAMB231 cells with control and GJB3 shRNAs. Interestingly, GJB3 knockdown alone did not significantly affect cell growth compared to controls in MDAMB231 cells (**Figure 2G**). However, when treated with DAC, MDAMB231 cells displayed a significant decrease in cell growth (**Figures 2H**). Similar observations were noticed in T47D cells (**Figure 2I, J and K**). These observations suggest that GJB3 expression might act as a resistance mechanism against DAC therapy in patients.

### 3.3 GJB3 inhibition creates cellular stress associated signaling and autophagy

Next to understand how GJB3 functions, we examined changes in gene expression after silencing it. We found 475 genes with increased expression and 801 genes with decreased expression following GJB3 knockdown (**Figure 3A).** Gene ontology analysis of these genes revealed that the upregulated genes were predominantly associated with stress-related pathways, including responses to starvation, and amino acid transport and synthesis processes (**Figure 3B and C**). Other significantly enriched pathways included the induction of autophagy and apoptosis. In contrast, downregulated genes were linked to cancer pathways.

**Figure 3:**
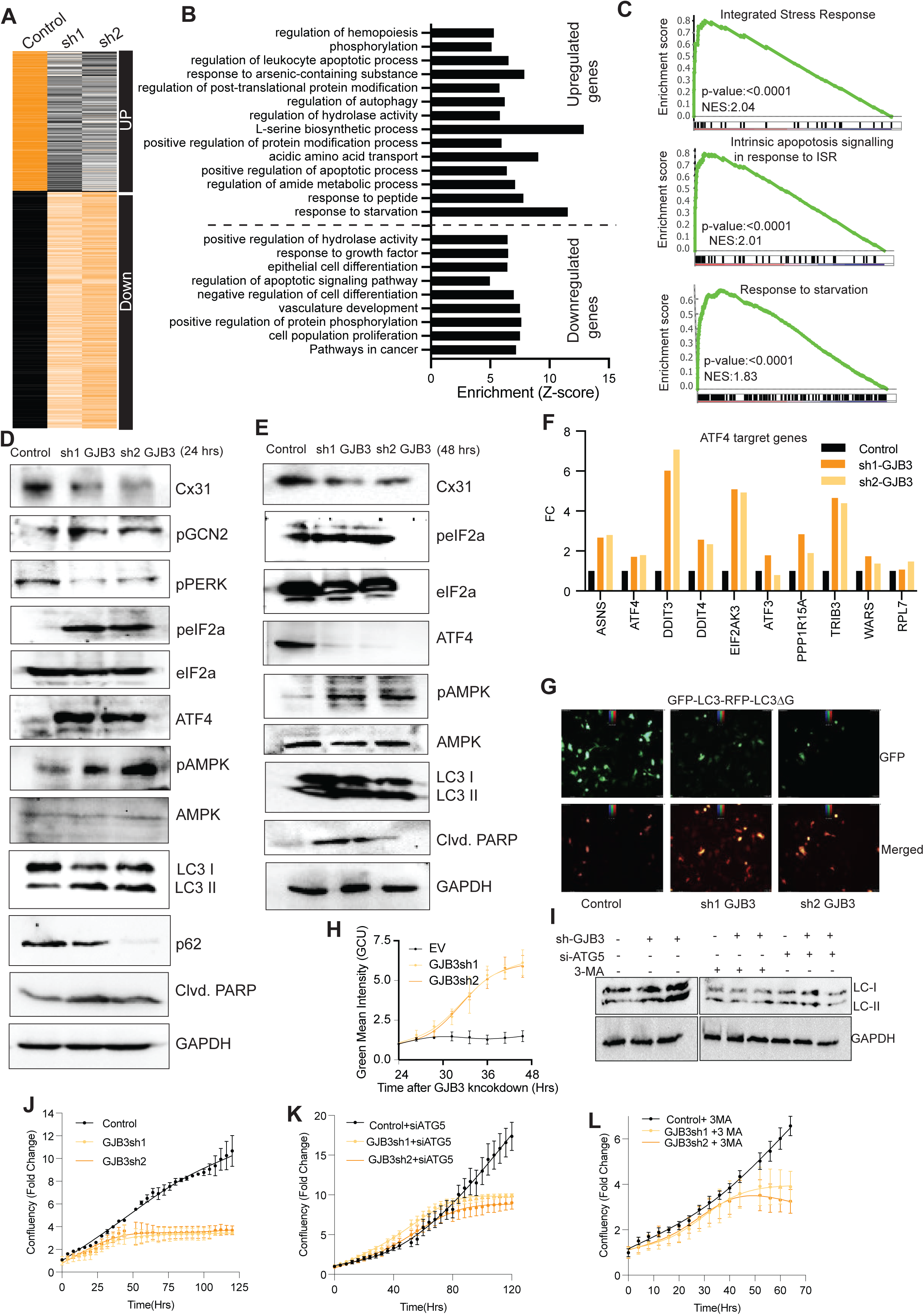
GJB3 Inhibition Induces Cellular Stress, Autophagy, and Apoptosis. **A)** SW480 cells with GJB3 knockdown were analyzed by RNA sequencing. Differentially expressed genes (upregulated and downregulated) were identified and their normalized expression values (Z-score transformed Transcripts Per Million - TPM) are visualized as a heatmap. **B)** Gene ontology (GO) analysis was performed using Metascape software on the differentially expressed genes identified in GJB3 knockdown cells. The most significantly enriched GO terms are presented. **C)** GO analysis was performed specifically on the upregulated genes identified in GJB3 knockdown cells. The top enriched terms are displayed in a bubble plot. **D)** Protein lysates were isolated from control and GJB3 knockdown SW480 cells after 24 hrs of knockdown. Western blotting was performed to determine the expression levels of specific proteins of interest. **E)** Protein lysates were isolated from control and GJB3 knockdown SW480 cells after 48 hrs of knockdown. Western blotting was performed to determine the expression levels of specific proteins of interest. **F)** Expression level of ATF target as determine by RNA-seq data in SW480 cells after treatment with control or shGJB3 plasmids. **G)** SW480 cells stably expressing both GFP-LC3 and RFP-LC3ΔG were used to assess the effect of GJB3 knockdown on autophagy. Cells were transfected with two different shRNAs targeting GJB3 or a control vector. Knockdown efficiency was confirmed, and then cells were imaged for GFP and RFP fluorescence. **H)** SW480 cells were used for GJB3 knockdown using control and GJB3 shRNAs. After knockdown the cells were stained with a caspase-3 activity detection reagent, and fluorescence intensity was measured at given time points. The line plot shown to detect the level of apoptosis of different cells. **I)** Control and GJB3 knockdown SW480 cells were transfected with siATG5 or treated with 3-methyladenine (3-MA), an inhibitor of autophagy. Protein lysates were then extracted and analyzed by Western blotting to assess LC3 protein levels. **J)** Control and GJB3 knockdown SW480 cells were plated, and their proliferation was monitored using an Incucyte live-cell imaging system. **K)** Control and GJB3 knockdown SW480 cells were transfected with siATG5, and then plated for proliferation analysis using the Incucyte live-cell imaging system. **L)** Control and GJB3 knockdown SW480 cells were treated with 3-MA and plated. Proliferation was measured using the Incucyte live-cell imaging system.

These findings from the GO analysis suggest that GJB3 knockdown triggers cellular stress, activating stress response pathways while inhibiting growth-related pathways. To validate this observation, we examined the stress response pathway in GJB3 knockdown cells compared to controls. Stress response pathway involves activation of eIF2α followed by ATF4 and has four sensors, GCN2, PERK, HRI and PKR (25, 26). First, we checked phosphorylation of eIF2α and found that GJB3 knockdown samples showed increased p-eIF2α levels and ATF4 level, suggesting activation of stress pathway in response to GJB3 knockdown (**Figure 3D**). eIF2α can be activated by four eIF2α kinases namely GCN2, HRI, PERK and PKR (25). In these GCN2 and PERK sense the amino acid starvation and ER stress (27). We checked the activation status of GCN2 and PERK and found increased phosphorylation of GCN2 but not PERK, suggesting that GJB3 knockdown induces amino acid stress (**Figure 3D**). Increase in p-eIF2α was sustained at later time point as well (**Figure 3E**). We also checked the expression of ATF4 targets and found consistent increase in ATF4 targets (**Figure 3F**). These observations suggest activation of GCN2-eIF2α-ATF4 pathway in response to GJB3 knockdown.

Since, GCN2 mediated ATF4 activation is an activator of autophagy (28). Also, pathway analysis had shown that GJB3 knockdown increase genes associate with positive regulation of autophagy (**Figure 3B**), we investigated the autophagy response in GJB3 knockdown cells. Interestingly, GJB3 knockdown led to increased levels of pAMPK and LC3II, alongside decreased p62 levels, suggesting a robust induction of autophagy flux (**Figure 3D and E**). This observation was further confirmed by measuring the ratio of GFP-LC3 to RFP-LC3ΔG (**Figure 3G**). It’s well-established that sustained stress-induced autophagy can trigger cell death via apoptosis (29). We checked the apoptosis level early and late time points after GJB3 knockdown. Interestingly, autophagy was induced after 24 hrs of GJB3 knockdown, we did not notice appreciable change in apoptosis activity as checked by PARP cleavage, caspase3/7 activity measurement (**Figure 3D and H**). However, there was robust apoptosis activation at later time points as measured by PARP cleavage and caspase3/7 activity (**Figure 3D and H** & **Supplementary Figure 2A**). We have also noticed proteasomal degradation of ATF4 at 48 hrs of GJB3 knockdown. ATF4 degradation is also a proapoptotic mechanism **(Figure 3D and H**). Further, to confirm if continuous autophagic activity is required for the cell proliferation defect in GJB3 knockdown cells, we treated GJB3 knockdown cells either with siATG5 or with 3-MA (**Supplementary figure 2B**). We showed that ATG5 inhibition or 3-MA treatment effectively blocked GJB3 induced autophagy (**Figure 3I**). The proliferation measurement of these cells showed that autophagy inhibition in GJB3 knockdown cells was able to partially rescue the proliferation defect. (**Figure 3J, K and L**). These findings suggest that GJB3 deficiency triggers cellular stress, leading to autophagy induction. Furthermore, sustained stress appears to drive autophagy-mediated apoptosis induction and cell death.

### 3.4 GJB3 knockdown causes cystine starvation in SLC7A11 low expressing cells

To further investigate metabolic changes induced by GJB3 knockdown, we performed mass spectrometry on methanolic extracts of control and knockdown cells. This analysis revealed alterations in numerous metabolites. Specifically, we observed decreases in L-Alanine, L-Cysteine, and L-Aspartate, as well as an increase in L-Proline levels in GJB3 knockdown cells (**Figure 4A and B**). We also leveraged Dependency Map data to categorize cells into GJB3-dependent and -independent groups. We then compared amino acid transporter expression between these groups using CCLE expression data. Notably, GJB3-dependent cells exhibited decreased expression of multiple amino acid transporters, while no transporters were upregulated (**Figure 4C**). Intriguingly, two of these downregulated transporters, SLC6A20 (Proline) and SLC7A11 (Cystine-Glutamate), were upregulated in GJB3 knockdown cells, as observed in microarray data (**Figure 4D**). The upregulation of SLC7A11 is associated with cystine starvation. Further analysis of mass spectrometry data showed that various cystine metabolites derivatives were also down regulated in GJB3 knockdown samples (**Figure 4E**). If GJB3 knockdown induces a cystine starvation-like phenotype, it should exhibit characteristic changes such as decreased glutathione (GSH) levels, reduced GSH activity, and increased lipid peroxidation (**Figure 4F**). We checked the GSH activity in GJB3 knockdown cells and found that GJB3 knockdown cells have lower GSH activity (**Figure 4G**). Further, we also showed that GJB3 knockdown cells have high lipid peroxidation level (**Figure 4H and I**). These observations suggest that GJB3 is associated with cystine transport.

**Figure 4:**
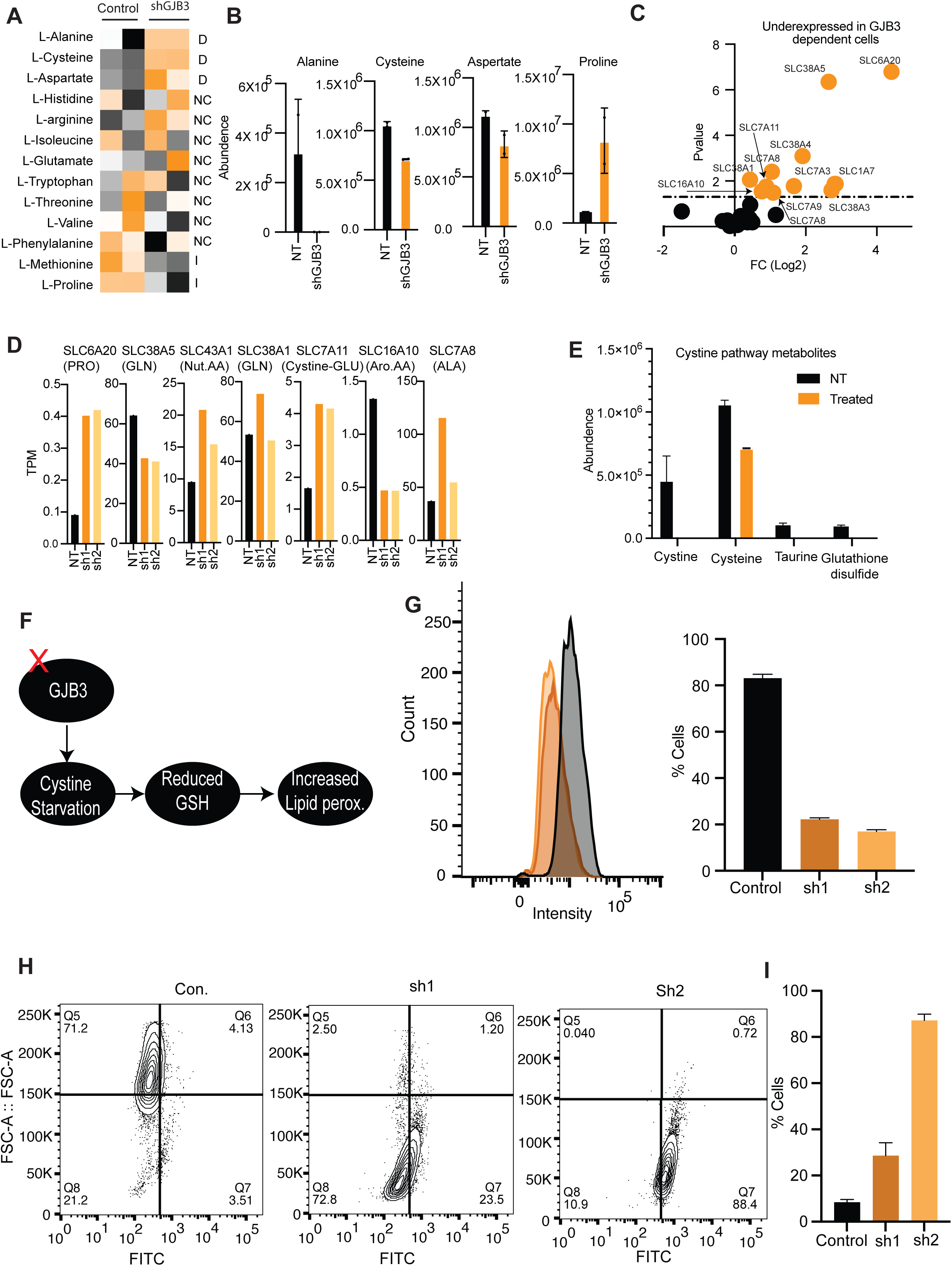
GJB3 is involved in Cystine transport. **A)** The methanol extract of Control and shGJB3 were subjected to metabolic profiling using mass-spectrometry. The abundance of detected amino acids are plotted in form heatmap. **B)** Amino acids whose abundance was changed in metabolomic analysis are plotted. **C)** Cell lines analyzed in Depmap were divided into GJB3 dependent and independent cells and expression of all the amino acid transporters were compared using t-test and a volcano plot was created. **D)** Expression of different amino acid transporters were analyzed from RNA-seq data and plotted. **E)** Abundance of metabolic derivatives of Cystine are plotted. **F)** A flowchart showing the proposed effect of GJB3 knockdown in GSH activity and lipid peroxidation. **G)** Control and GJB3 knockdown cells were tested for the GSH activity and analyzed using Flow cytometry. The results are plotted in histogram. The percent cell in each histogram was qualitied and plotted. **H)** Control and GJB3 knockdown cells were treated with Bodipy dye. The cells were anlysed using Flow cytometry and data is shown as contour plots. **I)** Control and GJB3 knockdown cells were treated with Bodipy dye. The cells were anlysed using Flow cytometry and quantified data is shown in bar plot.

Furthermore, we found that SLC7A11 expression was significantly higher in GJB3-independent cells compared to GJB3-dependent cells (**Figure 5A**). When we stratified lung and colon cancer patients with low GJB3 expression (associated with better survival) into high and low SLC7A11 groups, we observed that higher SLC7A11 expression correlated with poorer survival (**Figure 5B**). These findings suggest that GJB3 may play a role in cystine transport, particularly in cells with low SLC7A11 expression. To validate this hypothesis, we selected HT29 cells, which express high levels of both GJB3 and SLC7A11 (**Figure 5C**). Knocking down GJB3 in these cells did not significantly affect cell proliferation or long-term colony formation, suggesting that GJB3’s role in cystine transport may be more pronounced in cells with lower SLC7A11 expression (**Figure 5D, E and F**).

**Figure 5:**
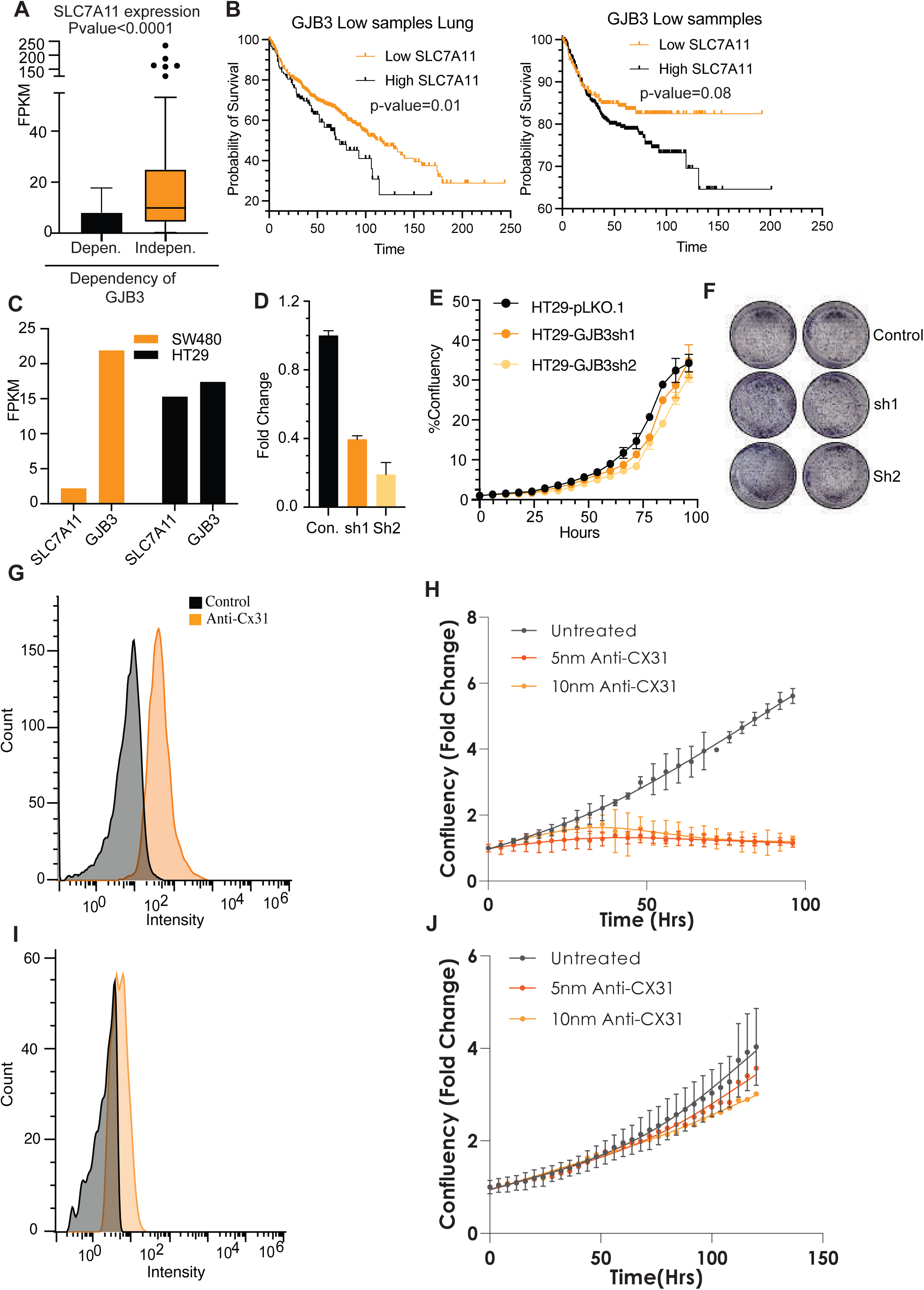
GJB3 cells compensate the SLC7A11 mediated cystine transport. **A)** Expression of SLC7A11 was compared between GJB3 depended and independent cells and plotted. **B)** Lung cancer and colorectal cancer patients were divided into GJB3 low and high expressing groups. GJB3 low expressing samples (which had good survival, figure 2 C and D) were further divided into SLC7A11 high and low samples and survival was compared. **C)** Expression of GJB3 and SLC7A11 in SW480 and HT29 cells are shown. **D)** HT29 cells were infected with control and GJB3 shRNA plasmids and expression of GJB3 was measured. **E)** HT29 cells were infected with control and GJB3 shRNA plasmids and proliferation of all the three cells were measured using incucyte. **F)** HT29 cells were infected with control and GJB3 shRNA plasmids and colony formation assay was performed for 15 days. Colonies were stained and photographed. **G)** SW480 cells were treated with antibody against GJB3 and stained with secondary antibody. The samples were analyzed for the surface binding of the antibody. **H)** SW480 cells were treated with antibody against GJB3 and incubated. Cell proliferation was measured using incucyte. **I)** HT29 cells were treated with antibody against GJB3 and stained with secondary antibody. The samples were analyzed for the surface binding of the antibody. **J)** HT29 cells were treated with antibody against GJB3 and incubated. Cell proliferation was measured using incucyte.

Further, to examine if GJB3 can be a therapeutic target, we checked the effect of GJB3 antibody on SW480 cells. We incubated the SW480 cells with GJB3 antibody and confirmed the binding of the antibody with Flow cytometry. The analysis confirms that GJB3 antibodies are specifically bound to Cx31 protein (**Figure 5G**). Next, we measure the effect of GJB3 antibody on SW480 proliferation. We noticed robust decrease in cell proliferation of SW480 cells in presence of SW480 antibody (**Figure 5H**). HT29 cells, which also express high level of Cx31 protein, did not show the effect of antibody similar to knockdown phenotypes (**Figure 5I and J**).

These observations suggest that GJB3 is involve in cystine transport and complement the SLC7A11 in cystine transport.

## 4. Discussion

Gap junctions are integral membrane proteins that facilitate intercellular communication, playing a pivotal role in maintaining tissue homeostasis and regulating tumor behavior (30). These structures are crucial in the process of carcinogenesis, as they allow for the direct transfer of ions and small molecules between adjacent cells, influencing various cellular processes (4). Despite their significance, most research has focused primarily on connexin 43 (Cx43), encoded by the GJA1 gene. Our work expands the understanding of gap junction proteins by analyzing genetic and epigenetic data from four prevalent adenocarcinomas to explore the regulation of all gap junction proteins. We have analyzed the genetic and epigenetic data from four most common adenocarcinomas to understand the regulation of all the Gap junction proteins. Our analysis identified GJB3 as one of the most interesting GAP junction genes in lung (LUAD) and colorectal cancer (COAD). Our study provides compelling evidence for the role of GJB3 in cancer progression and its potential as a therapeutic target. We have demonstrated that GJB3 is differentially regulated by DNA methylation in various cancer types, particularly in COAD and LUAD. In these cancers, hypomethylation of the GJB3 promoter leads to increased expression, which correlates with poor patient prognosis. Mechanistically, our findings suggest that GJB3 plays a critical role in maintaining cellular redox balance and amino acid metabolism. GJB3 knockdown induces cellular stress, leading to the activation of the integrated stress response (ISR) and autophagy. Notably, GJB3 appears to be particularly important for cystine uptake in cells with low SLC7A11 expression. This suggests that GJB3 may contribute to the metabolic reprogramming of cancer cells, allowing them to adapt to nutrient-limiting conditions. Our results highlight the potential of targeting GJB3 as a therapeutic strategy.

The initial analysis revealed that GJB3 is hypomethylated in lung adenocarcinoma (LUAD) and colorectal cancer (COAD), while exhibiting hypermethylation in breast cancer (BRCA) and prostate cancer (PRAD). Correspondingly, GJB3 expression levels were lower in BRCA and PRAD but higher in LUAD and COAD. Additionally, treatment with De-azacytidine (DAC) demonstrated that GJB3 expression is inhibited in BRCA and PRAD cells. Both the expression and methylation levels of GJB3 were predictive of patient survival outcomes: low levels in LUAD and COAD were associated with poor survival, whereas higher levels in BRCA and PRAD correlated with better survival rates.

These observations indicate that GJB3 may have distinct functional roles across different cancer types. Recent studies have also reported that GJB3 is downregulated in bladder cancer, which correlates with reduced cellular migration and invasion (23). Interestingly, our analysis showed that GJB3 is overexpressed in bladder cancer compared to normal bladder tissue, suggesting a complex role for GJB3 that may vary significantly depending on the specific cancer type and its microenvironment. The conflicting findings regarding GJB3’s expression highlight its context-dependent functionality.

Mechanistically, our research demonstrates that GJB3 knockdown induces cellular starvation, leading to the activation of the integrated stress response (ISR) pathway. Specifically, we found that this ISR activation is mediated through the phosphorylation of GCN2, a serine-threonine kinase that plays a critical role in sensing amino acid deficiency. GCN2 detects intracellular amino acid scarcity by binding to uncharged tRNAs, allowing it to respond to a general decline in amino acid availability rather than specific shortages (31). Our finding suggests that GJB3 is involved in amino acid transport, and its knockdown results in amino acid starvation. Amino acid starvation can trigger autophagy as a cellular survival response; however, if the starvation persists, it may also lead to apoptotic cell death (29). Our data indicate that within 24 hours of GJB3 knockdown, autophagy is activated without concurrent apoptosis. However, at later time points post-knockdown, apoptotic pathways become activated. We identified two key regulators of GJB3 knockdown-mediated autophagy. First, the activation of GCN2 due to amino acid starvation leads to the subsequent activation of ATF4, which upregulates autophagy-related genes such as SQSTM1. Indeed, we observed increased expression of several autophagy genes, including DDIT3, ERN1, SQSTM1, and TOM1, in cells with GJB3 knockdown. The second mechanism involves the activation of AMP-activated protein kinase (AMPK). AMPK is activated by decreased ATP levels and by the stress response gene Sestrin2 (32,33). Our analysis revealed both reduced ATP levels and elevated Sestrin2 levels following GJB3 knockdown, resulting in a robust increase in AMPK phosphorylation. Furthermore, we have shown that the activation of autophagy contributes at least partially to cell death associated with GJB3 knockdown. These findings highlight the intricate relationship between GJB3 expression, amino acid transport, and cellular stress responses, suggesting potential therapeutic avenues for targeting these pathways in cancer treatment.

Using mass spectrometry analysis, we investigated the levels of various amino acids and metabolites in GJB3 knockdown cells. Our findings revealed that cystine is one of the major amino acids whose levels are significantly affected by GJB3 knockdown. Additionally, we observed a decrease in other by-products of cystine metabolism following GJB3 knockdown. Cystine is typically transported into cells via the SLC7A11 antiporter. Analysis of DepMap data indicated that SLC7A11 expression is low in GJB3-dependent cells, and we found that cells with higher SLC7A11 expression were not adversely affected by GJB3 knockdown. These observations suggest that GJB3 is essential for amino acid transport, particularly cystine, in cells with low SLC7A11 expression.

Furthermore, we utilized a GJB3-specific antibody to demonstrate that GJB3 can be targeted in cell lines that depend on it for cystine uptake. This targeting could represent a promising therapeutic option for cancer cells that rely on GJB3.

While our study provides valuable insights into the role of GJB3 in cancer, it primarily focuses on colorectal and lung adenocarcinomas. Further research is needed to explore the role of GJB3 in other cancer types.

In conclusion, our findings suggest that GJB3 is a crucial regulator of cancer cell metabolism and survival. Targeting GJB3 may offer a novel therapeutic strategy for cancer treatment, particularly in tumors characterized by low SLC7A11 expression and high dependence on cystine uptake.

## Material and methods

### Plasmids

Primers targeting GJB3 were designed using Primer3Plus (https://www.primer3plus.com). The shRNA sequences for GJB3 knockdown were generated via the GPP Web Portal (https://portals.broadinstitute.org/gpp/public/) and subsequently cloned into the lentiviral vector pLKO.1 puro (Addgene, 8453). To study autophagy in GJB3-silenced cells, the autophagosome reporter plasmid DEST-CMV mCherry-GFP-LC3B WT (Addgene, 123230) and the autophagic flux reporter plasmid pMRX-IP-GFP-LC3-RFP-LC3ΔG (Addgene, 84572) were employed. Apoptosis detection was performed using the reporter construct pcDNA3-FlipGFP-T2A-mCherry (Addgene, 124429).

### Cell Culture

The cell lines used in this study were obtained from the American Type Culture Collection (ATCC) and maintained in the recommended growth medium supplemented with 10% fetal bovine serum (FBS) (Gibco, 10270106)) and 1% Penicillin-Streptomycin (PenStrep) (Gibco, 15140122) at 37°C in a 5% CO₂ environment. The Lenti-X 293T cell line (Takara, 632180) was utilized for lentiviral production through transfection using the Xfect Transfection Reagent (Takara, 631317), following the manufacturer’s protocol. Lentiviral infection was facilitated by polybrene-mediated transduction, and successfully infected cells were selected using puromycin (3 µg/mL).

For functional assays, in a 96-well plate, cell proliferation was monitored by plating 3,000 cells per well and measuring confluency over time using the Incucyte Live-Cell Imaging System. For the wound healing assay, 40,000 cells per well were seeded, and a scratch was introduced using the Incucyte WoundMaker. Colony formation assays were conducted by plating 1,000 cells per well, and after 14 days of incubation, colonies were fixed and stained with 0.25% Crystal Violet for visualization. For spheroid culture, cells were seeded onto Matrigel (Corning, 356234)-coated surfaces at a density of 0.3 million cells/mL. After a 30-minute incubation, 250 µL of Matrigel Matrix medium was added to the cell culture. The spheroids were allowed to develop for 14 days, after which they were imaged.

### Staining

Apoptosis was assessed using Caspase 3/7 Green Dye (Sartorius, 4440), and cell death was evaluated with Cytotox Red Dye (Sartorius, 4632); in both cases, cells were imaged and analyzed using the adherent Cell-by-Cell module of the Incucyte Live-Cell Analysis System.

### Treatments

To inhibit DNA methylation, cells were treated with 5-Aza-2′-deoxycytidine (10 µM; Sigma, A3656) for 96 hours. For autophagy inhibition, cells were exposed to 3-Methyladenine (5 mM; Sigma, M9281) for 24 hours. Knockdown of ATG5 was achieved by transfection with siATG5 using RNAiMAX reagent (Invitrogen, 13778075). To inhibit proteasomal degradation, cells were treated with MG132 (Sigma, 474790) at a concentration of 10 µM for 24 hours. To block Cx31, cells were incubated with the anti-Cx31 primary antibody (10 nM; CloudClone,PAB467Hu01) followed by detection with anti-rabbit IgG secondary antibody conjugated to AlexaFluor 647 (CST, #4414). These cells were subsequently analyzed using flow cytometry to quantify the binding.

### qRT-PCR

Total RNA was extracted by suspending cells in RNAiso Plus reagent (Takara, 9108) and isolating RNA using chloroform as per the manufacturer’s protocol. The purified RNA was quantified and reverse-transcribed into cDNA using the iScript cDNA Synthesis Kit (Bio-Rad, #1708841). For qRT-PCR, the synthesized cDNA served as a template in reactions prepared with Universal SYBR Green Mix (Bio-Rad, 172571), and TPT primers were included as internal controls. The data analysis was performed using the Bio-Rad CFX Maestro software.

### Western Blot

Proteins were extracted by suspending cell pellets in RIPA Lysis Buffer (G-Biosciences, 786-489) supplemented with 1× Protease Inhibitor Cocktail (Thermo, 87787). Following lysis, the samples were centrifuged at 13,000 × g for 12 minutes at 4°C, and the supernatant containing soluble proteins was collected. Protein concentrations were determined using the BCA Protein Assay Kit(Pierce, 23227). A total of 40 µg of protein from each sample was resolved on a 12% polyacrylamide gel via SDS-PAGE and subsequently transferred to a PVDF membrane (Bio-Rad, 1620177) using a wet transfer system. The membrane was blocked with 5% blocking powder (G-Biosciences 786-011) to prevent nonspecific binding and incubated with the appropriate primary antibodies overnight at 4°C.

After thorough washing, the membrane was incubated with the corresponding secondary antibodies for 90 minutes at room temperature. Following additional washes, the membranes were visualized using the FemtoLucent Plus HRP detection system (G-Biosciences, 786-003), and protein bands were imaged for analysis.

### Flow Cytometry

To analyze the cell cycle profile, cells were stained with Krishan Buffer containing propidium iodide (PI)(SRL, 25535-16-4) and incubated in the dark for 10 minutes. Stained cells were then analyzed using an Attune Flow Cytometer. Cellular glutathione levels were measured using the Cellular Glutathione Detection Assay Kit (CST, #13859) following the manufacturer’s protocol. Apoptosis was assessed using the Annexin V-FITC/PI Apoptosis Detection Kit (Elabscience, E-CK-A211) following the manufacturer’s instructions. Lipid peroxidation associated with ferroptosis was assessed using the BODIPY 581/591-C11 dye(Thermo, D3861). Cells were incubated with 0.5 µM BODIPY-C11 for 30 minutes. Following incubation, the cells were trypsinized, washed, and collected for analysis using a flow cytometer to quantify fluorescence intensity. All analyses were performed by Flow Jo(Version 10.9.0.).

### Scrape Loading/Dye Transfer Assay(SL/DT)

To evaluate gap junctional intercellular communication (GJIC) among cells, a confluent monolayer of cells was prepared and scraped using a sterile scalpel to create a wound. The well was then washed thoroughly with PBS to remove debris. Cells were incubated with Lucifer Yellow CH (1 mg/mL)(Thermo, L453) for 5 minutes to allow dye transfer through gap junctions. For the control group, cells were treated with Texas Red (1 mg/mL)(Thermo, D1828) under identical conditions. Following incubation, the cells were washed extensively with PBS to remove excess dye and imaged using a fluorescence microscope to assess dye transfer and localization. The extent of dye transfer was quantified using ImageJ and expressed as the Fraction of Control (FOC) using the formula:

GJIC FOC Treatment = (AreaTreatment(LY)-AreaTreantment(TR))/(AreaControl(LY)-AreaControl(TR)).

Here, LY represents the area stained by Lucifer Yellow, and TR denotes the area stained by Texas Red, with adjustments made for both the treated and control conditions.

### Metabolite Extraction

Control and experimental cells were seeded at a density of 0.7 million cells/mL in T25 flasks. After 24 hours, 1 mL of ice-cold 80% methanol (MeOH) was added to the flasks and incubated on ice for 5 minutes to arrest cellular metabolism. Cells were then harvested by scraping, transferred to 1.5 mL microcentrifuge tubes, and centrifuged at 13000 RPM for 30 minutes at 4°C. The resulting supernatant, containing the extracted metabolites, was carefully transferred into fresh tubes and analyzed via untargeted metabolomics using LC-MS/MS (Triple Quartz Metabolite Capture) for metabolite profiling.

## Acknowledgements

Sudhanshu Shukla would like to acknowledge Indian Council for medical research, Govt. of India (Grant # 2021-9513/CMB/ADHOC-BMS and EMDR/SG/13/2023-0244), Department of Biotechnology, Govt of India (Grant id – BT/PR51308/MED/30/2511/2023) and Anusandhan National Research foundation, Govt. Of India (Grant ID – CRG/2023/000837 for the funding. LLM based AI databases were used for the paraphrasing and Grammer correction.

## Author contributions

**DA:** Conceptualization, Methodology, Formal analysis, Investigation, review and editing.

**NB**: Investigation

**AC**: Investigation

**BKC**: Review and editing, Supervision

**SS**: Conceptualization, Methodology, Formal analysis, Writing – Review & Editing, Funding Acquisition.

---

**Supplementary figure 1:**
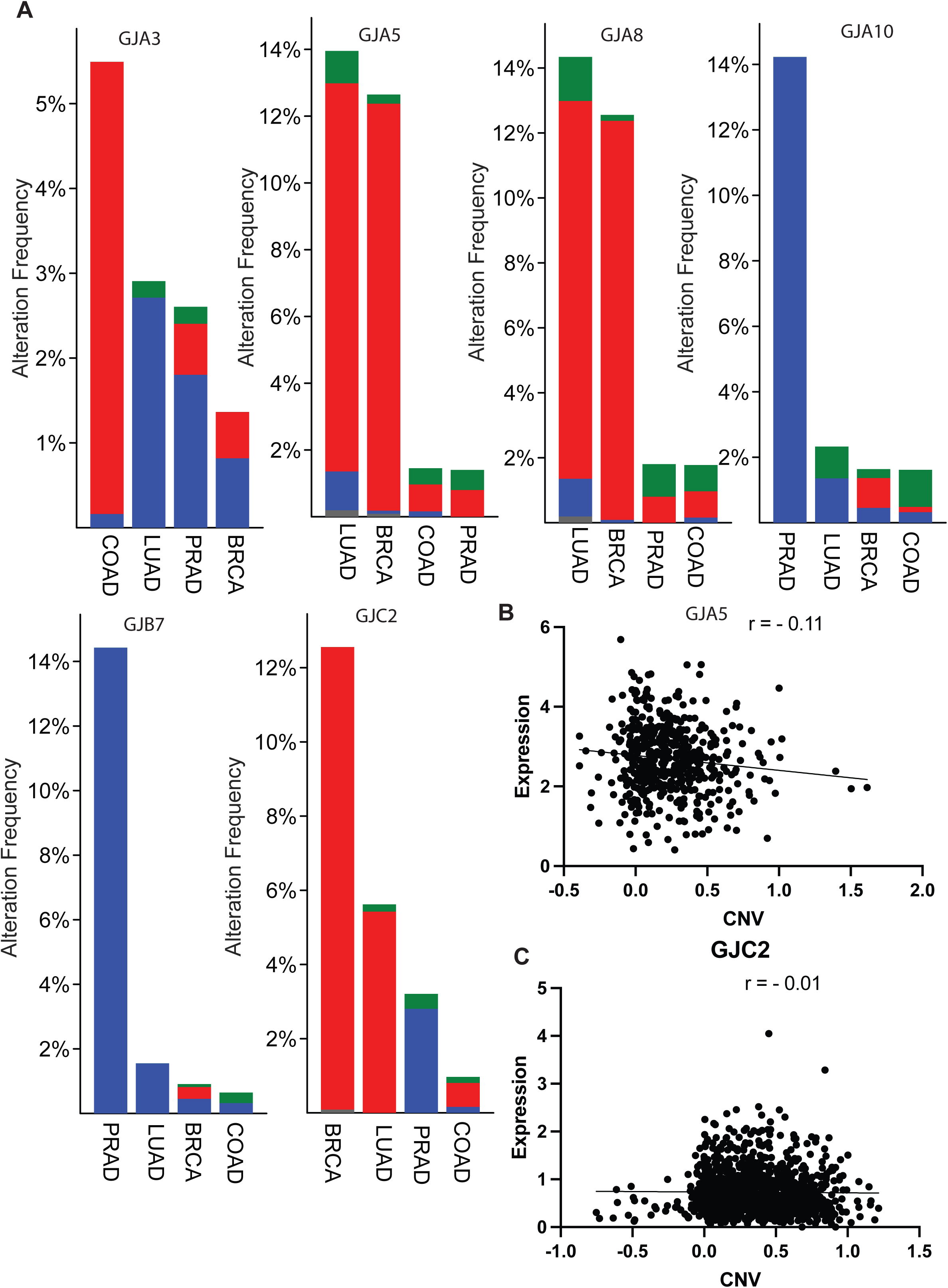
A) CNV and mutation frequency of mentioned GAp junction genes in four common adenocarcinomas B) Correlation between Expresion and CNV of GJA5 gene in LUAD samples. C) Correlation between Expresion and CNV of GJA5 gene in BRCA samples.

**Supplementarfy Figure 2:**
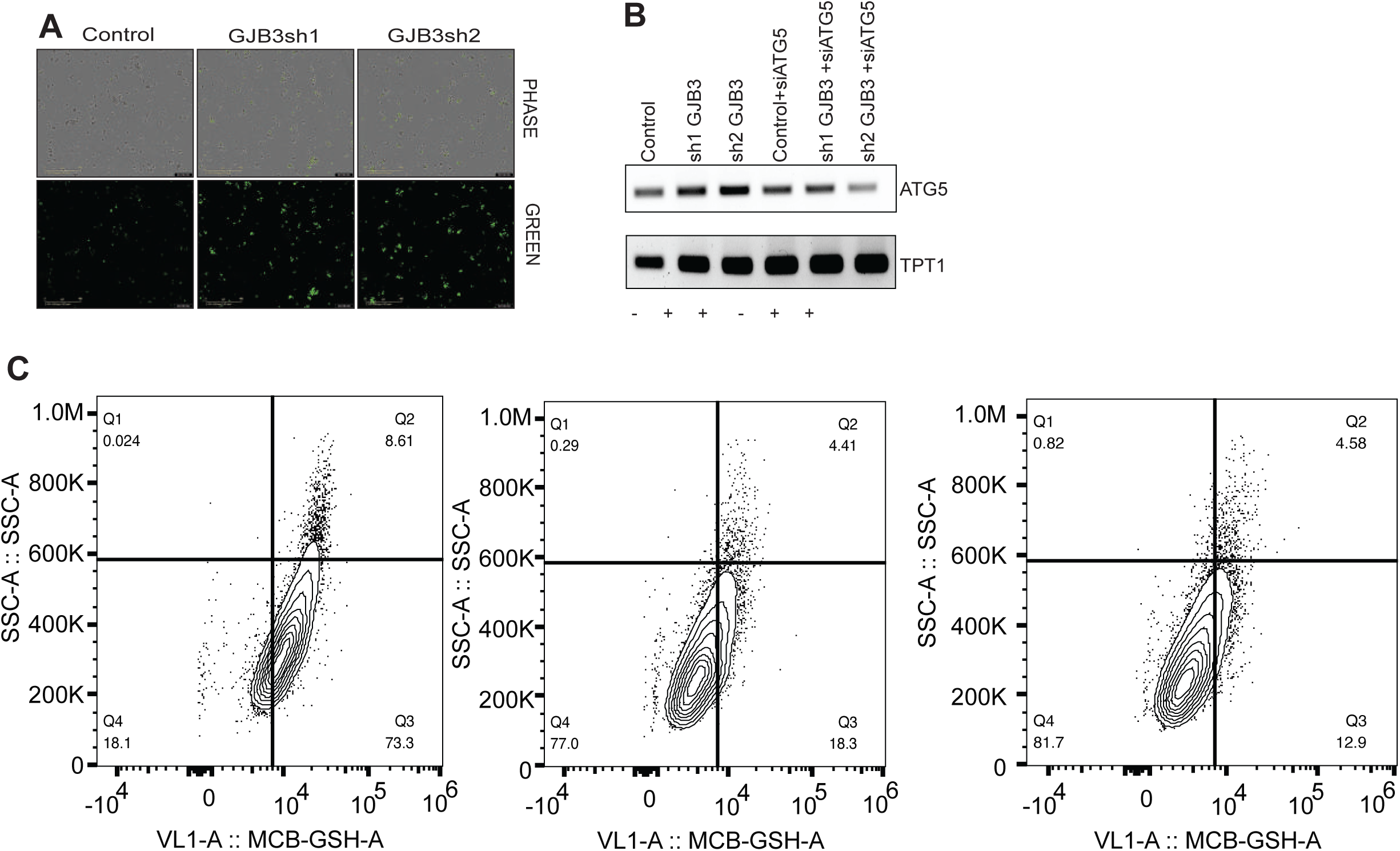
A) SW480 cells were used for GJB3 knockdown using control and GJB3 shRNAs. After knockdown the cells were stained with a caspase-3 activity detection reagent, and fluorescence intensity was measured using incucyte system. B) Control and GJB3 knockdown SW480 cells were transfected with siATG5. ATG5 expression levels were measured using semi-quantitative PCR followed by agarose gel electrophoresis. TPT1 was used as a control gene for normalization. C) Control and GJB3 knockdown cells were tested for the GSH activity and analyzed using Flow cytometry. The results are plotted in form of dot plot.

